# Adaptive Appraisals of Imaginary Objects

**DOI:** 10.1101/2021.08.18.456814

**Authors:** Magda L. Dumitru, Gitte H. Joergensen

## Abstract

When individuals think of objects or visual scenes in the absence of perceptual stimuli, they picture them on a virtual canvas, also known as “the mind’s eye”. What is currently unknown is the extent to which appraisals of imaginary objects are adaptive to contextual factors. Here, we present evidence that the eyes project imaginary objects in peripersonal space, that wearing sunglasses interferes with their projection, and that wearing a bicycle cap triggers emotional compensatory responses. Moreover, we show that maintaining a specific egocentric distance in the mind’s eye generates poor scaling of object similarity to object size, possibly due to an ego–allocentric computational overload. As a result, object representations are often retrieved from long-term memory only partially, with optimal representations requiring a perfectly still egocentric framework. We conclude that, like visual perception, mental imagery is adaptive to contextual factors with two important differences, namely external image projections and poor image resolution under load. We argue that these differences result from imagery being fully dependent on conscious computations and less on automatic processes.

## Introduction

Mental imagery, which refers to information processing in the ‘mind’s eye’, relies on subjective representations that support language and thought (Block 1983; Finke 1989; Hume 1784; Russell 1921). Furthermore, the brain anticipates interactive scenarios with relevant stimuli by constructing body-anchored maps extending, for example in peripersonal space around upper and lower limbs (for an overview, see Di Pellegrino and Ladavas 2015). We may thus assume that the mind’s eye extends in peripersonal space, such that objects are represented most accurately when people imagine them within arm’s reach. We explore this hypothesis in the current paper and further consider the impact on imaginary object representations of lighting, emotion, and imaginary distance, which are contextual factors known to elicit adaptive responses in visual perception.

Visual perception and imagery share basic processing mechanisms. For example, studies using the ‘blank screen’ paradigm (Altmann 2004) have shown that, when participants hear words or sentences referring to previously-viewed objects, they move their gaze over an empty screen or in complete darkness to the location where these objects had stood (Ferreira, Apel, and Henderson 2008; Johansson, Holsanova, and holmkvist 2006; Laeng and Teodorescu 2002; Richardson and Spivey 2000). These findings demonstrate that working memory traces are maintained in the mind’s eye, which behaves like visual sensors by flexibly monitoring objects situated away from an observer. Incidentally, directing attention outwardly via eye movements is a key assumption of the earliest model of visual processing (aka ‘extramission’ - for a review, see Gross 1999), which remains popular among children and adults (Cottrell and Winer 1994; Winer and Cottrell 1996; Winer et al. 2002; Gregg et al. 2001) and further accounts for the belief that the eyes project some form of energy or invisible force onto others (i.e., scopaesthesia), possibly with evil intent (i.e., ‘casting the evil eye’).

Neuroscientific evidence points to a common substrate of imagery and perception (Kastner et al. 1998; Kosslyn, Ganis, and Thompson 2001; Pylyshyn 2002) and validates the dominant neuroscientific model of visual imagery (Pearson et al. 2015; Dijkstra, Bosch, and Van Gerven 2019), according to which the primary visual cortex and the ventral cortical visual stream contribute to identifying and classifying imaginary objects (Kosslyn, Thompson, and Ganis 2006; Pearson 2019). Importantly, visual stimuli also engage the dorsal visual stream, which is involved in object localization and potential interaction with the observer (Goodale and Milner 1992), although visual stimuli elicit activation in ventral areas even in settings that typically activate the dorsal stream, albeit only for nearby objects (Grill-Spector et al. 1999; Grill-Spector, Kourtzi, and Kanwisher 2001; Kourtzi and Kanwisher 2000; James et al. 2002; Vaziri-Pashkam and Xu 2017; Xu 2018). Moreover, research on primates has found that the ventral premotor cortex continued to be activated by an object placed in its field even after turning off the lights or removing the object (Graziano, Hu, and Gross 1997).

For imaginary objects, it is unclear how their appraisals accommodate changes in egocentric distance, which should recruit the dorsal visual stream in addition to the ventral stream that is typically involved in mental computation (Committeri et al. 2004; Hoshino et al. 2005). Current evidence supports a naturalistic relationship between size and similarity appraisals of imaginary objects. For example, when participants were asked to evaluate the size and similarity of objects of the same kind, which were mentioned but not seen, they rated their size on a scale that was inversely correlated with the scale of their similarity, such that the smaller the objects, the more similar to each other they appeared to be (Dumitru and Joergensen 2015). Thus, the apparent distance between objects in a group increases with object size, which allows for better discrimination of their properties and lowers similarity ratings. This inverse relationship between imaginary size and similarity corresponds to predictions by the ‘size-distance invariance hypothesis’, also known as the ‘size invariance hypothesis’ in visual processing – for a review, see Epstein, Park, and Casey 1961). According to this hypothesis, the farther away an observer stands from visual objects, the smaller these objects appear to be (Foley 1972; Gilinsky 1951), typically at a lower rate (aka “egocentric distance underestimation”) than warranted by physical distance (Loomis et al. 1992; Thomson 1983).

To evaluate the impact of contextual factors on imaginary object appraisals, we designed two studies, as follows. In the ‘distances’ study, we aimed to determine whether appraisals are adaptive to imagery-internal factors that is, to variations in imagined ego- and allocentric distance and their interaction. Participants were asked to imagine themselves standing close by (the CLOSE condition) or far away (the FAR condition) from imaginary objects. In the ‘headgear’ study, we aimed to determine whether appraisals are adaptive to imagery-external factors that is, lighting conditions (i.e., participants wore sunglasses in the SUN condition) and compensatory behavior afforded by safety gear (i.e., participants wore a bicycle cap in the CAP condition). The task in both studies was to evaluate the size or similarity of imaginary objects that were referred to by plural names (e.g., “rabbits”). We contrasted experimental conditions with a control condition (CTRL), where participants evaluated object size or similarity without further instructions or equipment.

If appraisals are adaptive to changes in egocentric distance, imaginary objects in the CLOSE and in the FAR condition should be rated as larger or smaller, respectively, than in the control condition. We might also find comparable size appraisals in one experimental condition and the control condition, which would indicate that the mind’s eye operates at a specific distance from the observer. Moreover, by comparing size appraisals in the control and the SUN conditions, where the luminosity of projections is filtered as they leave the eyes, thus lowering image resolution, we could establish whether images are indeed projected outside the body. In contrast, size appraisals in the control condition should not differ significantly from the CAP condition, as bicycle caps do not interfere with eye gaze. Furthermore, the size invariance hypothesis predicts good scaling, albeit inverse, between size appraisals and similarity appraisals, which are proxies for image resolution (i.e., higher or lower similarity ratings reflect lower or higher resolution respectively). This scaling is particularly relevant in the distances study, where egocentric distance could affect allocentric distance appraisals. In contrast, wearing sunglasses can only impact object properties retrieved from long-term memory, and wearing a bicycle cap can only trigger emotion-based responses that exclusively engage the ventral stream (40-41) to compensate for the associated sensation of increased safety (42).

## Methods and materials

### Participants

We recruited 360 volunteers for the distances study and 270 volunteers for the headgear study. All participants were students at the University of Connecticut, had English as their mother tongue, and had normal or corrected-to-normal vision. All participants signed an informed consent form upon enrollment, in accordance with the Declaration of Helsinki, and received course credit for testing sessions lasting up to 60 minutes. The protocol was approved by the Ethics Committee of the University of Connecticut. Volunteers were randomly assigned to one of six groups of 60 individuals each in the distances study, and to one of six groups of 45 individuals each in the headgear study. Participants in the first three groups in the distances study were asked to evaluate the size of objects in one of three egocentric distance conditions (close, far, and control). Participants in the remaining three groups were asked to evaluate the similarity of objects to each other in the same egocentric distance conditions. In the headgear study, participants in the first three groups were asked to evaluate the size of objects in one of three headgear conditions (sunglasses, cap, and control), whereas participants in the remaining three groups were asked to evaluate object similarity in the same conditions.

### Stimuli

We selected word stimuli (43) referring to animals, plants, and tools (30 × 3 = 90 words) in the distances study and to animals, plants, tools, and vehicles (111 + 41 + 85 + 37 = 274 words) in the headgear study. All words were individually randomized and presented in plural form (e.g., ‘bees’, ‘apples’, ‘forks’, ‘bicycles’) written onscreen, one word per trial, accompanied by a visually-presented rating scale.

### Design and procedure

Volunteers were seated at a comfortable distance in front of a computer screen and reacted to each plural word they saw (e.g., ‘rabbits’) by moving a cursor horizontally on a scale from 0 to 10. In the distances study, participants in the first group were instructed to imagine themselves positioned close-by (i.e., a couple of feet away) from the objects referred to by each word and evaluate them from ‘not very big’ (0) to ‘very big’ (10). Participants in the second group were instructed to imagine themselves positioned far away (i.e. ‘tens of feet away’) from the objects referred to by each word and rate object size on the same scale. Participants in the third group were not provided with any information on their position with respect to the objects, but were simply instructed to rate their size on the 0-to-10 scale. Participants in the fourth, fifth, and sixth groups rated the objects referred to by words for their similarity to each other (e.g., how similar rabbits in a group would be from each other) on a scale from ‘0” (i.e. not very similar to each other) to “10” (i.e., very similar to each other) while imagining themselves standing close-by, far-away, or not positioned at any particular distance from the objects. In summary, the distances study followed a 3 (Object position: CLOSE versus FAR versus Control) x 2 (Evaluation: Size versus Similarity) between-subjects factorial design.

In the headgear study, participants in the first three groups were instructed to evaluate object size while wearing sunglasses, a bicycle cap, or no headgear at all respectively. The remaining three groups were instructed to rate object similarity in the same headgear conditions. For both studies, the dependent variable was the score between 0 and 10 given to each word stimulus. We compiled average scores for each item and for each condition prior to statistical analysis. In summary, the second study followed a 2 (Headgear: SUN versus CAP versus Control) x 2 (Evaluation: Size vs. Similarity) between-subjects factorial design.

## Results

### Average object size and similarity appraisals

Figure 1 shows mean ratings for size and similarity in the distances study and in the headgear study. In a series of *t*-tests, we determined that, in the distances study, object size ratings were comparable in the CTRL and the CLOSE conditions, *t* (178) = 1.036, *p* = 0.301, but objects projected FAR were smaller than those in the CTRL condition, *t* (178) = 3.17, *p* = 0.001, Cohen’s *d* = 0.473. However, object appraisals were less similar to each other in the CTRL condition than in either the CLOSE condition, *t* (178) = 12.64, *p* < 0.001, Cohen’s *d* = 1.88 or the FAR condition, *t* (178) = 13.27, *p* < 0.001, Cohen’s *d* = 1.97.

**Figure 1.**
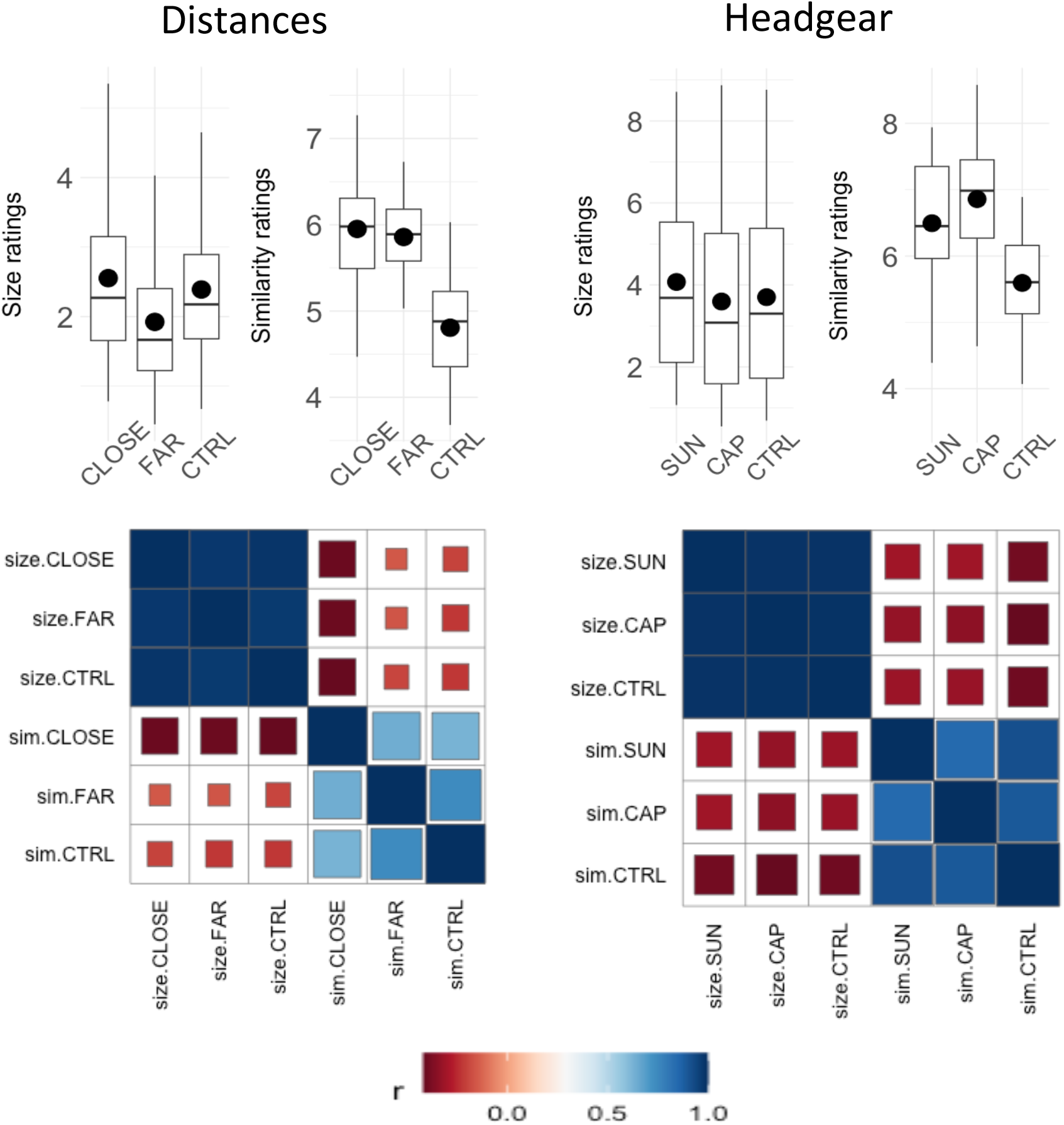
Average ratings of imaginary objects. In the distances study, CTRL size ratings were comparable with CLOSE ratings, but higher than FAR ratings. In the headgear study, CTRL size ratings were comparable with CAP ratings but lower than SUN ratings. For object similarity, CTRL ratings were higher than those for experimental conditions in both studies. Black dots indicate average values. Size and similarity appraisals were inversely correlated in both studies, supporting the size invariance hypothesis.

In the headgear study, objects were larger in the SUN condition than in the CTRL condition, *t* (546) = 2.59, *p* = 0.009, Cohen’s *d* = 0.22, but comparable in the CAP and the CTRL conditions, *t* (546) = 0.370, *p* = 0.711. Moreover, objects in the CTRL condition were rated as being less similar to each other than in either the SUN condition, *t* (546) = 13.38, *p* < 0.001, Cohen’s *d* = 1.14 or the CAP condition, *t* (546) = 17.76, p < 0.001, Cohen’s *d* = 1.51.

Correlation matrices between size ratings and similarity ratings for objects situated close, far, and CTRL followed the size invariance hypothesis and confirmed previous findings (Dumitru and Joergensen, 2016) that imaginary object size and similarity appraisals are inversely correlated.

### Object size and similarity appraisals across categories

For each study, we built correlation matrices for size and similarity ratings across categories. For the headgear study, we randomly selected a subset of the data to obtain an equal number of items across categories (30 × 4 = 120 words) before building correlation matrices. As shown in Figure 2, size ratings were robust for categories across conditions in both studies, with intra-category Pearson correlation coefficients close to 1 and p-values well below 0.05. Similarity ratings were slightly less robust, as within-category correlations were lower compared to size correlations.

**Figure 2.**
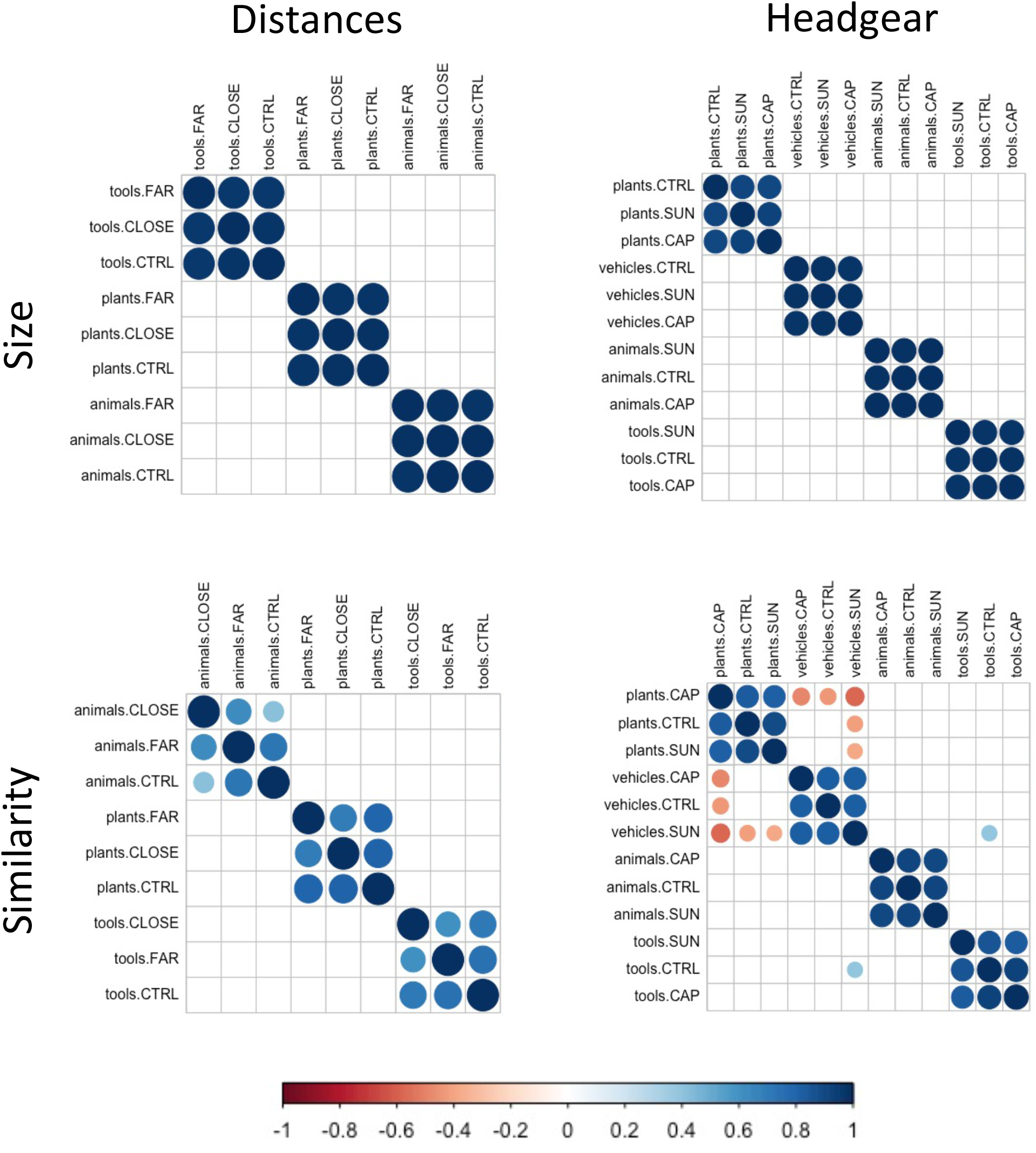
Size and similarity ratings of imaginary objects by categories. Size and similarity ratings of imaginary objects across categories. In both studies, size ratings were more robust than similarity ratings. Correlation matrices display significant Pearson correlations (p < 0.05) between object categories and suggest good retrieval of object categories from long-term memory.

### Object size and similarity appraisals across individual stimuli

We used Mantel tests (Mantel 1967) across dissimilarity matrices that we computed separately for individual ratings of stimuli size and similarity. Unlike correlation tests (e.g., Pearson’s), which evaluate linear relationships between two variables (e.g., size ratings in the CLOSE vs. the control condition), the Mantel test can reveal patterns across different types of data (i.e., size and similarity ratings across CLOSE, FAR, and CTRL conditions). As seen in Figure 3, the two dissimilarity matrices, one for size and the other for similarity ratings, are visually very different in the distances study, but comparable in the headgear study. Indeed, the Mantel test was non-significant in the distances study (*r* = 0.043, *p* = 0.184), but significant in the headgear study (*r* = 0.127, *p* = 0.001), suggesting that the scaling of similarity ratings to size ratings across conditions was different in the distances study, but not in the headgear study.

**Figure 3.**
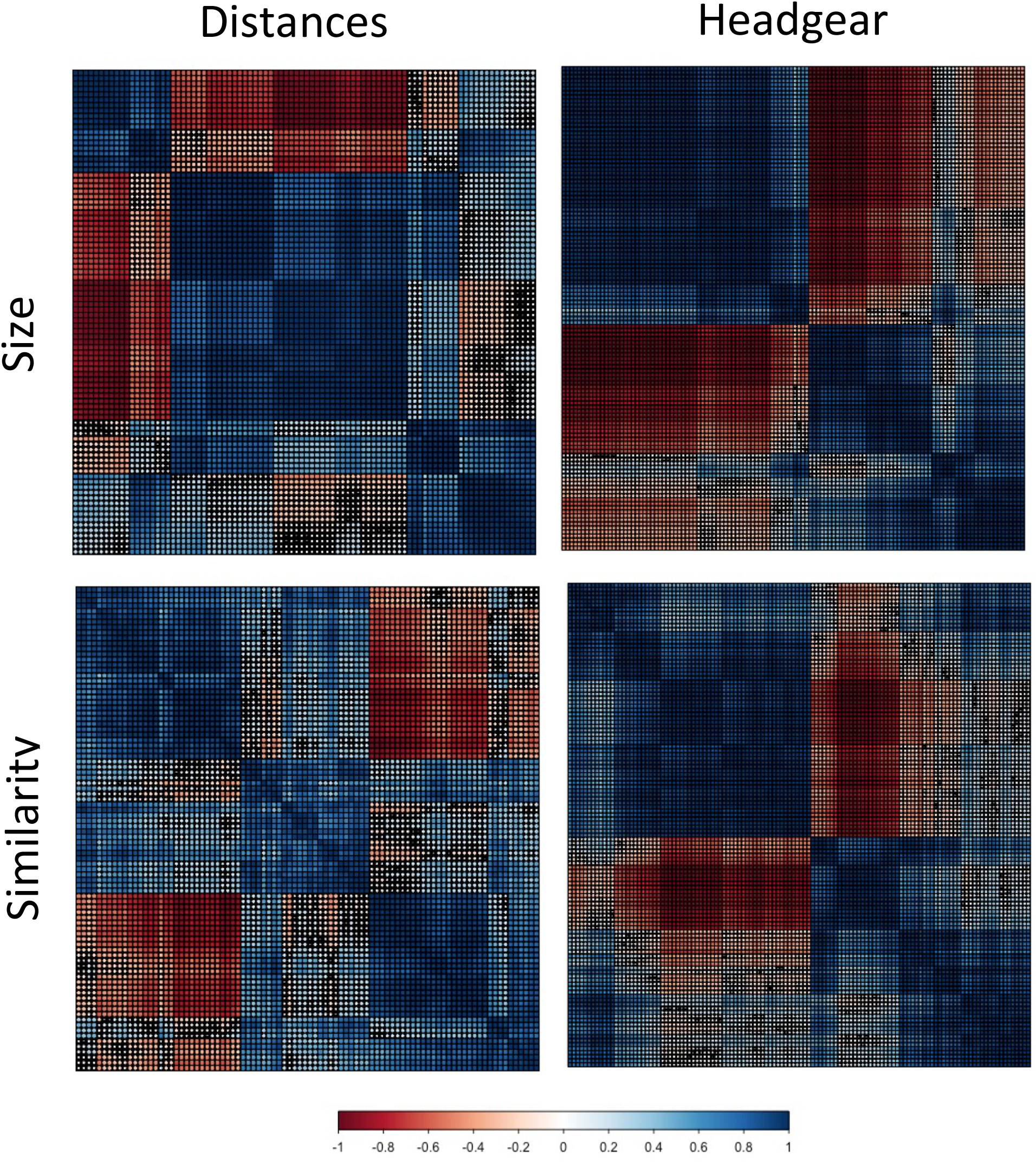
Size and similarity ratings of imaginary objects by items. Size and similarity ratings of imaginary objects across individual stimuli. Dissimilarity matrices were computed separately for size and similarity. Mantel tests over size and similarity matrices were significant in the headgear study, but non-significant in the distances study, suggesting an ego-allocentric computational overload (NB. Featured values are for the 120 items used in category analyses, but results were comparable over all items).

## Discussion

### Adaptation to peripersonal space and light

In the distances study, we found a parallel between vision and mental imagery regarding the orienting of attention outside the body (Spivey and Geng 2001; Posner 1980), such that object size appraisals were comparable when participants were asked to imagine objects situated close by (i.e., a couple of feet away) and when they received no egocentric distance instructions (i.e., the control condition). These findings demonstrate that the default distance between an observer and imaginary objects is relatively short, hence the mind’s eye extends outside the body in peripersonal space, which is typically within arm’s reach (Rizzolatti et al. 1981). We sought further support for this hypothesis by asking participants to wear sunglasses in the headgear study. We found that size appraisals adapted to lower light, which interfered with imaginary object projections. Thus, objects appeared larger than in the control condition. We may account for this difference by assuming flexible manipulation of the appraised objects that the observer could bring closer, or by a shift in imaginary egocentric framework for better inspection of the imagined objects. Nevertheless, higher size ratings did not result in better allocentric estimations (i.e., smaller similarity ratings). We may account for these findings by a drop in resolution for imaginary objects, afforded by poor lighting. Overall, these findings reveal that the eyes are instrumental for projecting imaginary objects outwardly, which is compatible with the extramission hypothesis (cf. Gross 1999).

### Adaptation to emotion

As part of the headgear study, we asked participants to wear a bicycle cap while rating imaginary objects for size or similarity. The words referring to these objects were not chosen to signal imminent danger, for which participants would need to develop coping strategies (Adams and Hillman 2001). However, merely wearing safety gear was found to be sufficient for triggering compensatory responses to visual stimuli (Dumitru and Pasqualotto 2018). Emotional contexts are known to influence object representation via the ventral visual stream (Kryklywy and Mitchell 2014, Ta, Liu, Brennan, and Enns 2010; Vuilleumier 2005), which connects structurally and functionally with the amygdala, where mainly negative emotions are being processed (Morris et al. 1998). We found that imaginary objects appeared comparable in size but more similar to each other in the CAP condition compared to the control condition, suggesting that lower image resolution could be due to less attention being assigned to imaginary object representations in emotion-laden contexts. Importantly, these adaptive processes are likely to engage exclusively the ventral visual stream and suggest incomplete retrieval from long-term memory of object representations given poor allocentric estimation (i.e. similarity ratings) compared to the control condition. Overall, our findings suggest that imaginary object appraisals adapt to emotional contexts.

### Adaptation to egocentric and allocentric distance

In the distances study, we explored the impact of imaginary egocentric distance manipulations on imaginary object appraisals, including adaptation to concurrent egocentric and allocentric distance computations in similarity ratings. We found size appraisals of imaginary objects to be adaptive to egocentric distance, such that objects imagined to be far from an observer were rated as being smaller than objects imagined to be at the default distance, as predicted by the size invariance hypothesis (Epstein, Park, and Casey 1961). Moreover, as predicted by the size invariance hypothesis, imaginary objects appeared more similar to each other in the FAR condition than in the control condition, but they did not appear less similar to each other in the CLOSE condition. Thus, similarity ratings in both the CLOSE and the FAR conditions were higher than in the control condition, suggesting poor allocentric distance estimation when participants were asked to concurrently observe a specific egocentric distance from imaginary objects.

Poor allocentric computations for CLOSE objects in the distances study could be due to a conflict between ego- and allocentric computations, which are supported by neuronal activity in the dorsal and the ventral visual stream respectively, although this distinction is not clear cut (Fairchild et al. 2025; Rolls 2024; Schenk and McIntosh 2010; Snyder et al. 1998). Indeed, neuroscientific evidence supports a partial division of labor when processing egocentric distance, with activation of dorsal areas alongside ventral areas for nearby but not for far-away visual stimuli (Grill-Spector et al. 1999; Kourtzi and Kanwisher 2000; James et al. 2002; Vaziri-Pashkam and Xu 2017; Xu 2018; Lacquaniti and Caminiti 1998). For mental imagery, we would need to assume a back-and-forth processing loop between allocentric and egocentric information in the ventral stream or across streams, in which case distinct computations in the FAR and the CLOSE conditions would underlie poor allocentric appraisals. However, we also found inverse correlations between size and similarity ratings in both studies, thus validating size-invariant computations in the mind’s eye that is, immediate scaling, however approximate, of similarity ratings to assumed object size. These findings suggest a common cause for poor similarity ratings in in the distances study.

Alternatively, the dorsal stream would encode both egocentric and allocentric information, with no role for ventral visual areas in visual perception (Snyder et al. 1998; Crowe, Averbeck, and Chafee 2008; Rizzolatti and Matelli 2003; Schenk 2006), we may posit an all-egocentric computation in the ventral stream for imaginary objects. However, this possibility is unlikely in our study, where participants received explicit instructions that ensured conscious retrieval of information from long-term memory. Indeed, we reminded participants in each trial of the experimental conditions (i.e., CLOSE and FAR) to imagine themselves standing either close or far from the objects. Thus, even for size appraisals, which typically rely on the dorsal stream in visual perception, we would need to assume considerable input from the ventral stream, whose content alone reaches visual awareness (59). Moreover, correlations between object size and similarity across categories were high in both studies irrespective of egocentric distance, emotion, or lighting, dovetailing with previous findings that ventral representations are impervious to contextual factors (Grill-Spector, Kourtzi, and Kanwisher 2001; Andersen 1997). Nevertheless, we did find significant differences between average similarity appraisals of control and experimental conditions in both studies, lending support to a weaker version of the size invariance hypothesis (e.g., Ayzenberg and Behrmann 2022).

This explanation would allow us to posit comparable mechanisms of similarity appraisals that is, equally imperfect representations for items in groups of imaginary objects in the FAR and in the CLOSE conditions. Assuming that both size and similarity appraisals of imaginary objects involve the ventral visual stream, possibly alongside the dorsal visual stream based on their joint involvement in mental processing (Committeri et al. 2004; Koshino et al. 2005), we may infer that size appraisals (i.e., egocentric computations) in the distances study are approximations of size-invariant objects, and that similarity appraisals (i.e., ego-allocentric computations) in the same study betray a computation overload. We sought further support for this explanation from Mantel tests, which we ran between size and similarity ratings across all conditions in each study. We found good scaling in the headgear study (test statistic was significant) but poor scaling in the distances study (test statistic was not significant), inviting the inference that imagery is a graded phenomenon. Thus, variable retrieval from long-term memory of a subset of object attributes (i.e., size and category) depending on context would be due to a computational overload of combined egocentric and allocentric information in ventral areas (for a review of the impact of long-term memory on ventral visual stream processes in visual perception see Rolls 2024; Martin and Barense 2023).

To summarize, we have shown that, like visual perception, mental imagery is adaptive to internal and external contextual factors including lighting, emotion, and egocentric distance. However, we also found particularities, which are likely to stem from the very nature of mental imagery, rather than from differences in processing. Thus, in the absence of external stimuli, images projected outside the body (aka extramission). Moreover, when participants consciously maintained a specific egocentric distance (CLOSE or FAR) throughout the task, they experienced an ego–allocentric computational overload resulting in failure to upload the full range of imaginary object properties when estimating object similarity. Our findings are also compatible with previous reports in the literature that shifts in egocentric framework interfere with mental imagery. Saccades, for instance, even though inherent to the proper functioning of the visual system, were shown to have a disruptive effect on mental imagery (Irwin and Brockmole 2000; Martin, Shao, and Boff 1993; Van Duren 1993). In our studies, changes in lighting or emotion as well as instructions to keep a specific distance from imagined objects increased object similarity, suggesting that optimal representation of imaginary objects require a perfectly still egocentric framework.

## Acknowledgments

The authors are grateful to study participants and to Gerry Altmann, who made it all possible.

## Author contributions

MLD: conceptualization; formal analysis; investigation; methodology; software; visualization; writing – original draft preparation. GHJ: data curation; methodology; software; validation; writing – review & editing.

## Data availability

The authors will provide access to datasets to interested individuals upon reasonable request.

